## Supporting Information for "Neurocognitive Dynamics of Preparatory and Adaptive Cognitive Control: Insights from Mass-Univariate and Multivariate Pattern Analysis of EEG data"

| **Table S1**  *ANOVA table for linear mixed effects regression model of reaction time.* | | | | | |
| --- | --- | --- | --- | --- | --- |
| Predictor | Sum of Squares | DF_num_ | DF_den_ | F-value | P-value |
| Age | 0.0025 | 1 | 71.69 | 0.0468 | 0.8294 |
| Sex | 0.3937 | 1 | 70.71 | 7.4263 | 0.0081 |
| Accuracy (Acc) | 7.4192 | 1 | 89.0 | 139.9624 | < 2.2e-16 |
| Cue | 0.2969 | 1 | 111.33 | 5.6007 | 0.0197 |
| Probe | 0.0340 | 1 | 703.97 | 0.6407 | 0.4237 |
| Acc x Cue | 8.3643 | 1 | 111.33 | 157.7914 | < 2.2e-16 |
| Acc x Probe | 4.7790 | 1 | 703.97 | 90.1542 | < 2.2e-16 |
| Cue x Probe | 0.0097 | 1 | 124.03 | 0.1831 | 0.6695 |
| Acc x Cue x Probe | 3.6634 | 1 | 124.03 | 69.1096 | 1.39e-13 |
| Note. num = numerator; den = denominator. Effect of covariates: Age did not significantly influence RT (p = 0.829, d = −0.02, 99% CI = [−0.30, 0.25]). In contrast, we observed a small but significant effect of gender on RT, with females responding slightly slower than males (∆MF −M = 0.035 seconds, 99% CI = [0.001, 0.069], d = 0.29, 99% CI = [0.01, 0.57], p = 0.006). | | | | | |

| **Table S2**    *ANOVA table for linear mixed effects regression model of error rates.* | | | | | |
| --- | --- | --- | --- | --- | --- |
| Predictor | Sum of Squares | DF_num_ | DF_den_ | F-value | P-value |
| Age | 1.4688 | 1 | 49 | 3.3292 | 0.0742 |
| Sex | 1.1416 | 1 | 49 | 2.5875 | 0.1141 |
| Cue | 10.7386 | 1 | 153 | 24.3403 | 2.085e-06 |
| Probe | 68.4619 | 1 | 153 | 155.1767 | < 2.2e-16 |
| Cue x Probe | 82.3628 | 1 | 153 | 186.6847 | < 2.2e-16 |
| Note. num = numerator; den = denominator. Effect of covariates: Age did not significantly influence the error rate (p = 0.070, d_low−high_ = 0.25, 99% CI = [−0.12, 0.63]). In contrast, the model estimated a small but significant effect of participants’ gender on the error rate, with females achieving slightly lower error rates than males (∆M_F −M_ = −0.011, 99% CI = [−0.031, 0.008], d = −0.22, 99% CI = [−0.59, 0.15], p = 0.109). | | | | | |

**Figure S1**

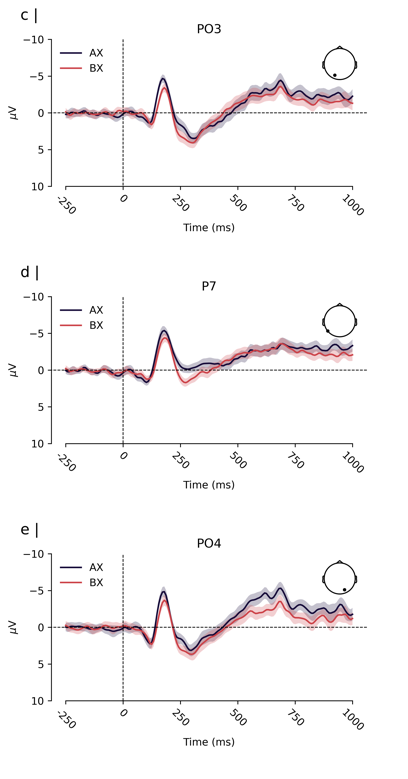

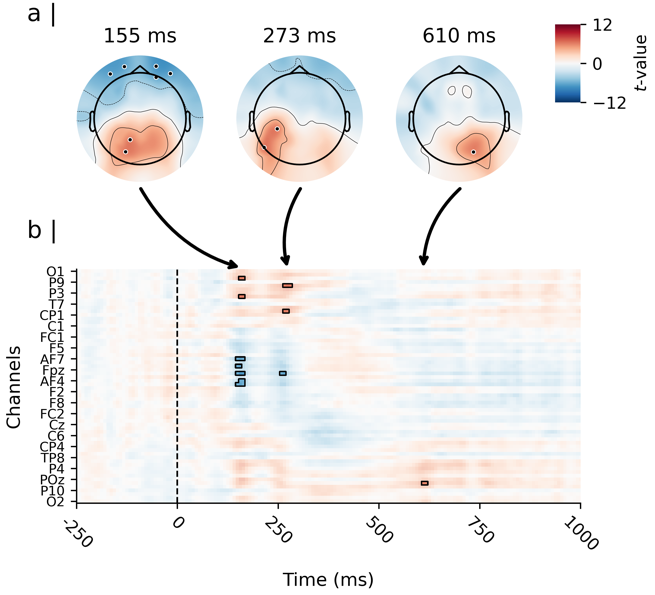

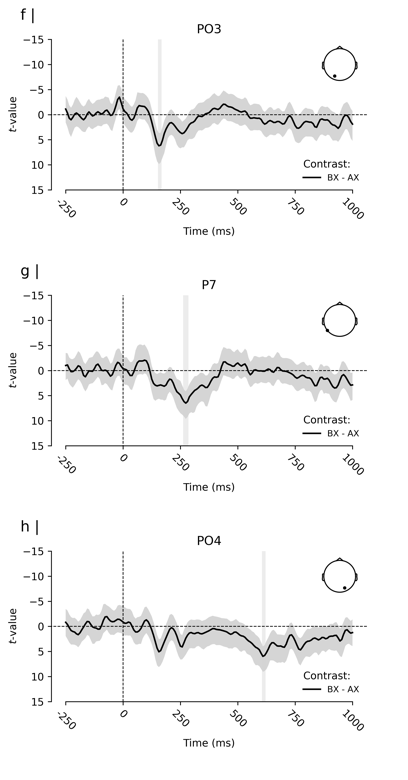
*Results of the mass univariate analysis of the probe-evoked amplitude response for the contrast “AX” minus “BX”*

*Note.* Depicted are the results of a paired-samples t-test contrasting “X” probes in “B” cue trials minus “X” probes in “A” cue trails (i.e., the contrast of probe evoked activity between “BX” and “AX” pairs, time locked to probe onset). Panel a) shows the topographical distribution of the estimated effect (i.e., the t-values) on the scalp for three time windows with the strongest effects. However, we found no significant differences and therefore no sensors are highlighted. Panel b) shows the estimated effects for the complete probe processing interval and the entire sensor space. Time is depicted on the X-axis in b), with axis-ticks appearing every 250 milliseconds (ms). Only every 4th sensor is labelled on the Y-axis in b) to avoid clutter. Panels c), d), and e) depict the Event-Related-Potential (ERP, i.e., the grand average of the participants’ mean evoked amplitude response) for both conditions (i.e., the average probe evoked activity by “BX” and “AX” pairs, time locked to probe onset). Depicted are sensors that were located within the topographical maps depicted in a). The solid lines depict the ERP for each condition. The coloured shades surrounding the lines express the uncertainty of the estimates with a 99% within-subjects confidence interval (cf. 73). Panels f), g), and h) depict the time course of the estimated difference effect at each corresponding sensor. The solid line depicts the t-value for the contrast “BX” - “AX” probe, and the shaded region surrounding the line expresses its uncertainty with a 99% confidence interval.

**Figure S2**

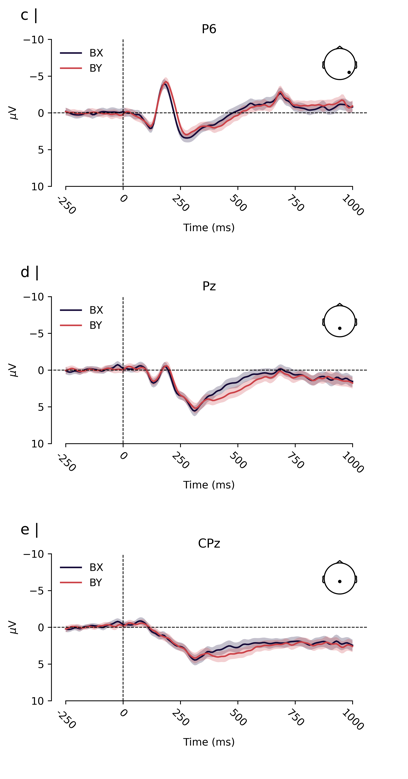

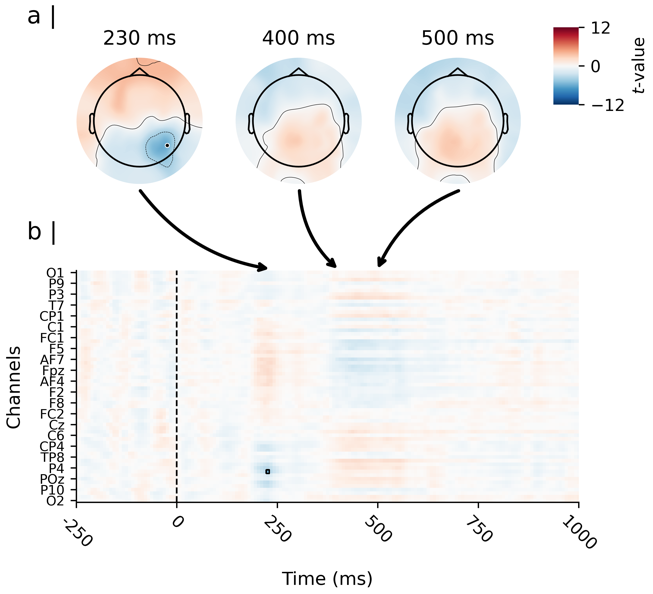

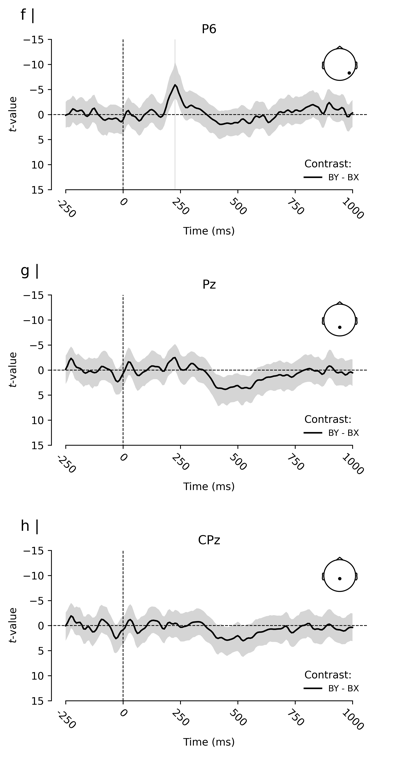
*Results of the mass univariate analysis of the probe-evoked amplitude response for the contrast BY minus BX*

*Note.* Depicted are the results of a paired-samples t-test contrasting “Y” probes in “B” cue trials minus “X” probes in “B” cue trails (i.e., the contrast of probe evoked activity between “BY” and “BX” pairs, time locked to probe onset. Panel a) shows the topographical distribution of the estimated effect (i.e., the t-values) on the scalp for three time windows with the strongest effects. However, we found no significant differences and therefore no sensors are highlighted. Panel b) shows the estimated effects for the complete probe processing interval and the entire sensor space. Time is depicted on the X-axis in b), with axis-ticks appearing every 250 milliseconds (ms). Only every 4th sensor is labelled on the Y-axis in b) to avoid clutter. Panels c), d), and e) depict the Event-Related-Potential (ERP, i.e., the grand average of the participants’ mean evoked amplitude response) for both conditions (i.e., the average probe evoked activity by “BX” and “BY” pairs, time locked to probe onset). Depicted are sensors that were located within the topographical maps depicted in a). The solid lines depict the ERP for each condition. The coloured shades surrounding the lines express the uncertainty of the estimates with a 99% within-subjects confidence interval (cf. 73). Panels f), g), and h) depict the time course of the estimated difference effect at each corresponding sensor. The solid line depicts the t-value for the contrast “BX” - “BY” probe, and the shaded region surrounding the line expresses its uncertainty with a 99% confidence interval.

**Figure S3**

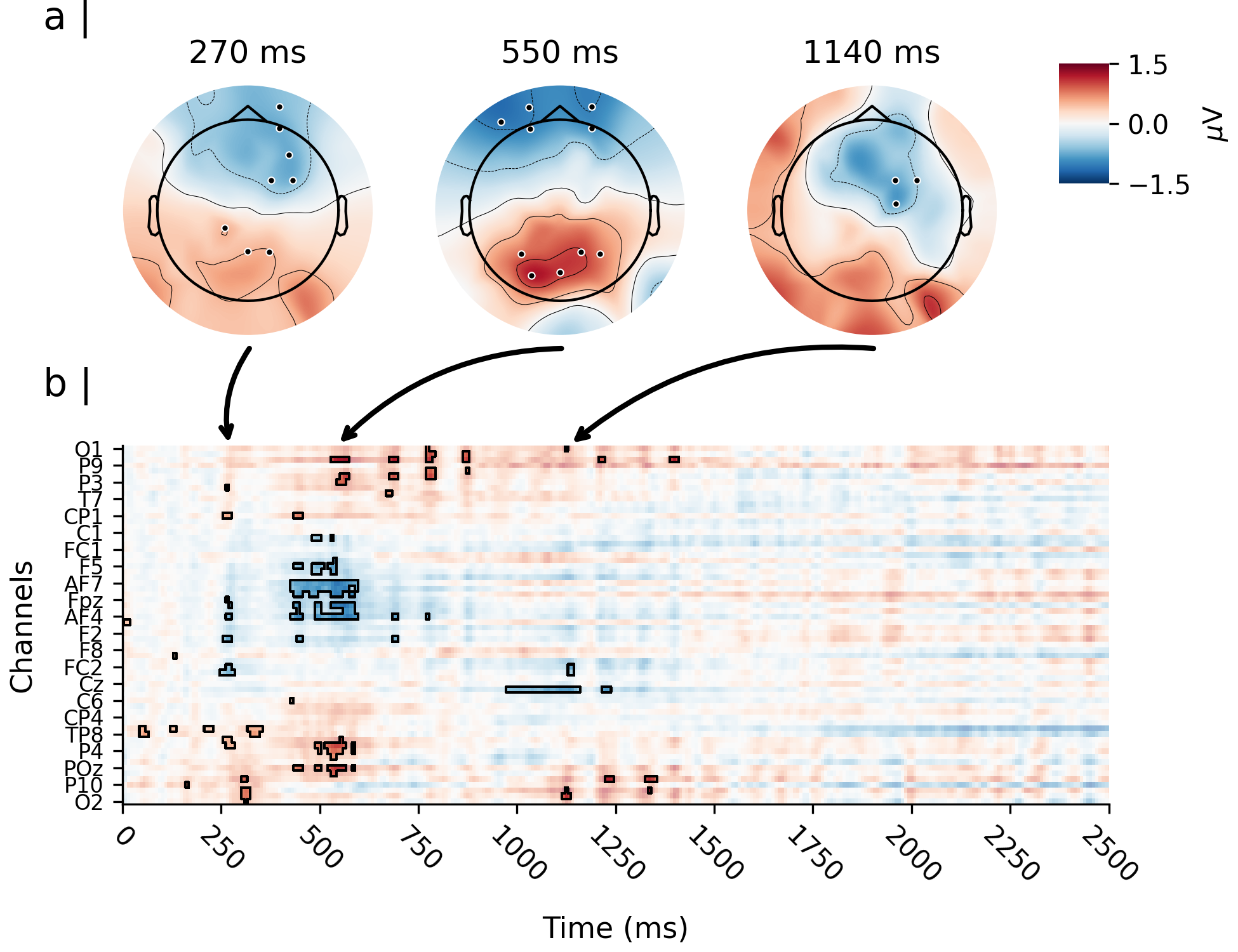
*Effect of d’ Context on the differential amplitude response between cues “B” - “A”*

Note. Depicted is the moderating effect of d’ Context on the estimated difference between cue “B” and cue “A”. Panel a) shows the topographical distribution of the estimated moderation effect (i.e., the regression weight of *d’context* predicting the difference between “B” and “A” cues) on the scalp for three representative spatio-temporal clusters. Significant channels (i.e., sensor for which the 99% CI of the moderation effect did not include zero) are highlighted. Panel b) shows the estimated effects for the complete cue retention interval and the entire sensor space, time locked to the onset of the cue. Time is depicted on the X-axis in b), with axis-ticks appearing every 250 milliseconds (ms). Only every 3rd sensor is labelled on the Y-axis in b) to avoid clutter. The highlighted time samples in b) are significant.

**Figure S4**

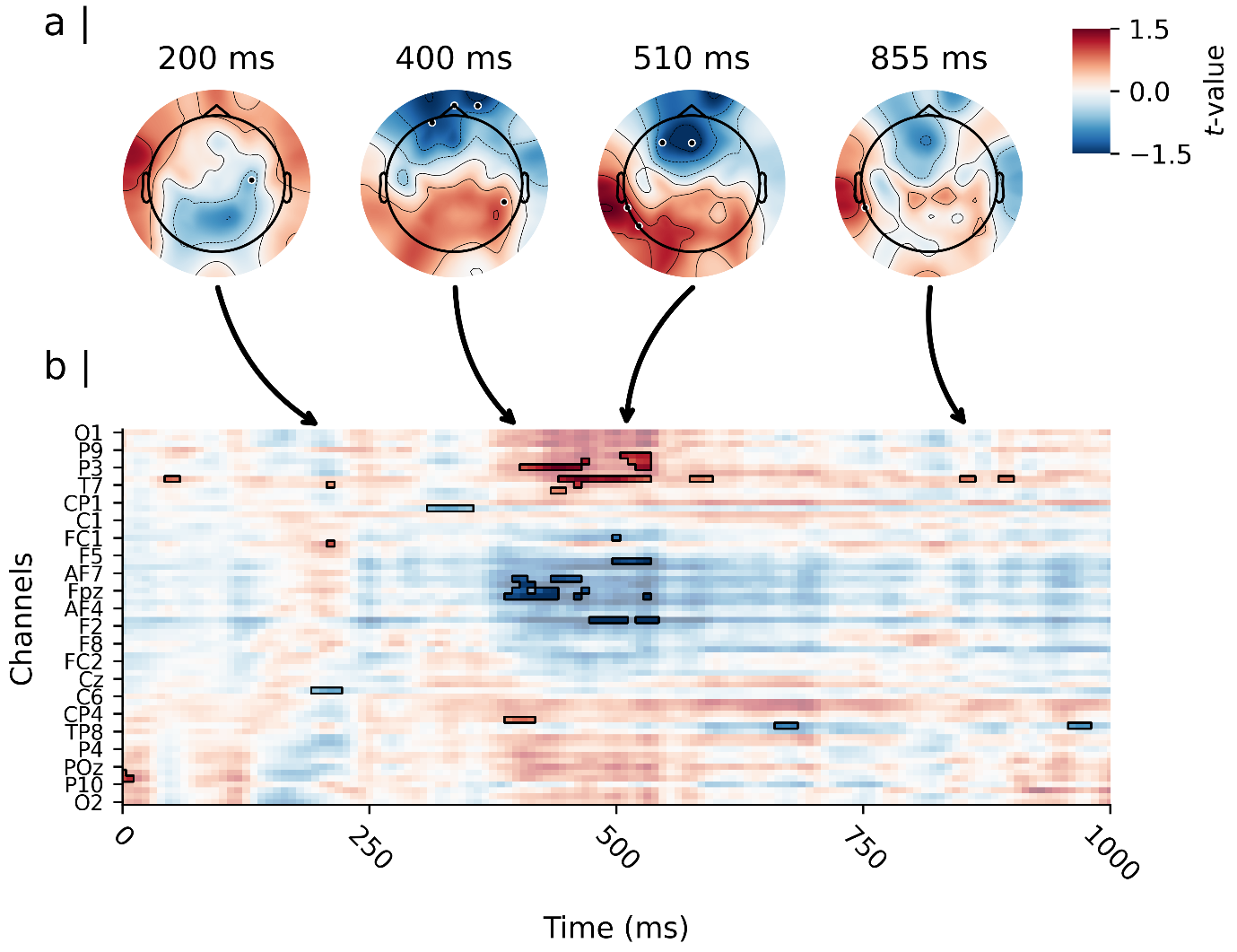
*Effect of d’ Context on the differential amplitude response between probes “Y” - “X” in “A” cue trials.*

Note. Depicted is the moderating effect of d’ Context on the estimated difference between probe “Y” and probe “X”, when presented after an “A” cue (i.e., the effect of d’ Context on the difference between “AY” and “AX” pairs, time locked to probe onset). Panel a) shows the topographical distribution of the estimated moderation effect (i.e., the regression weight of d’context predicting the difference between “AY” and “AX” trials) on the scalp for three representative spatio-temporal clusters. Significant channels (i.e., sensor for which the 99% CI of the moderation effect did not include zero) are highlighted. Panel b) shows the estimated effects for the complete cue retention interval and the entire sensor space, time locked to the onset of the cue. Time is depicted on the X-axis in b), with axis-ticks appearing every 250 milliseconds (ms). Only every 3rd sensor is labelled on the Y-axis in b) to avoid clutter. The highlighted time samples in b) are significant.

**Figure S5**

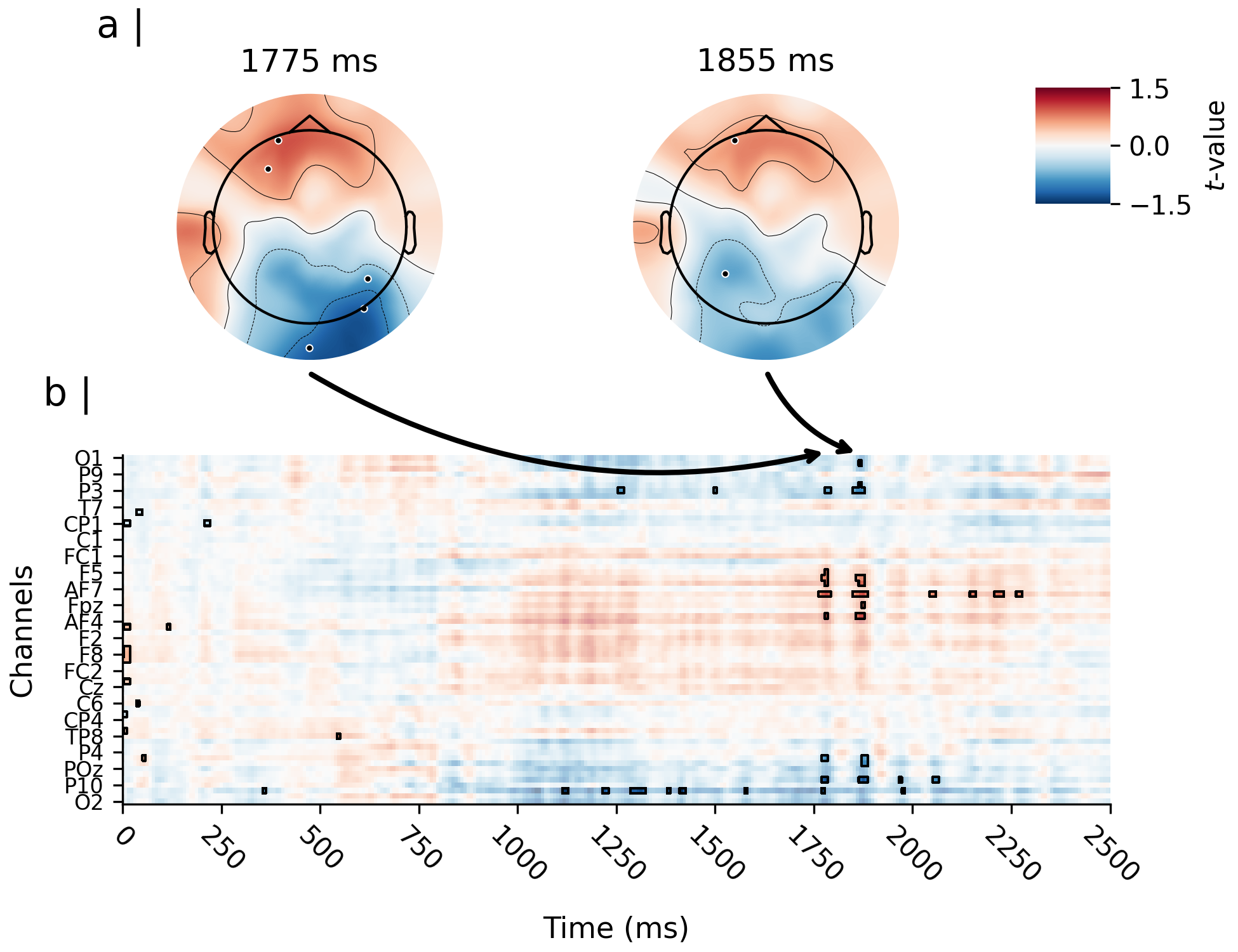
*Effect of A-cue Bias on the differential amplitude response between cues “B” - “A”.*

Note. Depicted is the moderating effect of the A-cue bias on the estimated difference between cue “B” and cue “A”. Panel a) shows the topographical distribution of the estimated moderation effect (i.e., the regression weight of A-cue bias predicting the difference between “B” and “A” cues) on the scalp for three representative spatio-temporal clusters. Significant channels (i.e., sensor for which the 99% CI of the moderation effect did not include zero) are highlighted. Panel b) shows the estimated effects for the complete cue retention interval and the entire sensor space, time locked to the onset of the cue. Time is depicted on the X-axis in b), with axis-ticks appearing every 250 milliseconds (ms). Only every 3rd sensor is labelled on the Y-axis in b) to avoid clutter. The highlighted time samples in b) are significant.

**Figure S6**

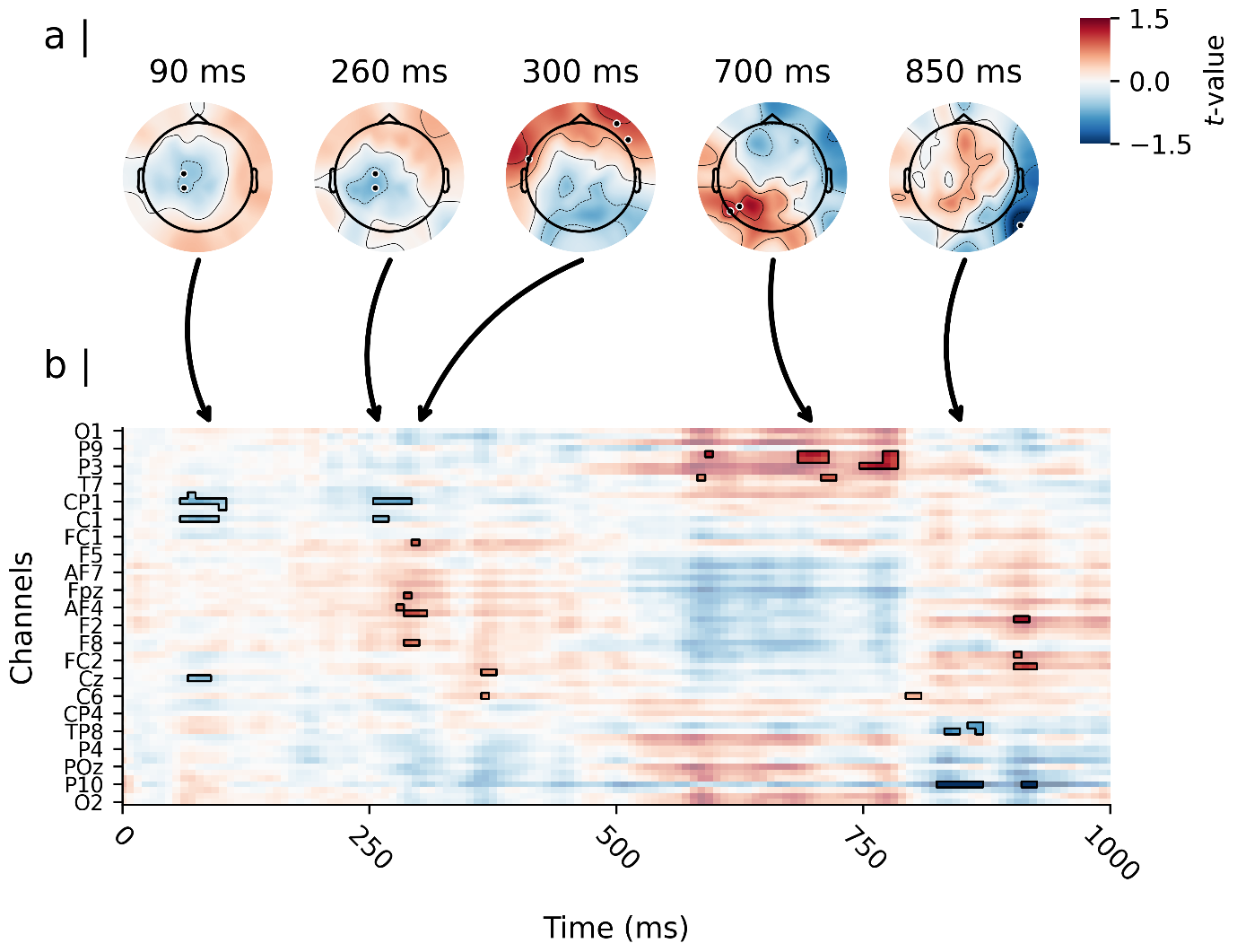
*Effect of A-cue bias on the differential amplitude response between probes “Y” - “X” in “A” cue trials.*

Note. Depicted is the moderating effect of the A-cue bias on the estimated difference between probe “Y” and probe “X”, when presented after an “A” cue (i.e., the effect of A-cue bias on the difference between “AY” and “AX” pairs, time locked to probe onset). Panel a) shows the topographical distribution of the estimated moderation effect (i.e., the regression weight of the A-cue bias predicting the difference between “AY” and “AX” trials) on the scalp for three representative spatio-temporal clusters. Significant channels (i.e., sensor for which the 99% CI of the moderation effect did not include zero) are highlighted. Panel b) shows the estimated effects for the complete cue retention interval and the entire sensor space, time locked to the onset of the cue. Time is depicted on the X-axis in b), with axis-ticks appearing every 250 milliseconds (ms). Only every 3rd sensor is labelled on the Y-axis in b) to avoid clutter. The highlighted time samples in b) are significant.
